# The soybean (*Glycine max* L.) cytokinin oxidase/dehydrogenase multigene family; identification of natural variations for altered cytokinin content and seed yield

**DOI:** 10.1101/2020.09.24.311738

**Authors:** Hai Ngoc Nguyen, Shrikaar Kambhampati, Anna Kisiala, Mark Seegobin, RJ Neil Emery

**Author notes:** **Declarations**. **Ethics approval:** Not applicable. **Consent to participate:** Not applicable. **Consent for publication:** Not applicable. **Availability of data and material:** The datasets generated during and/or analysed during the current study are available from the corresponding author on reasonable request. **Code availability:** Not applicable. **Author’s contributions**: HNN: Data curation, Writing - Reviewing and Editing; SK: Investigation, Data curation, Visualization, Writing - Original draft preparation, Writing - Reviewing and Editing; AK: Data curation, Writing - Reviewing and Editing; MS: Data curation, Writing - Reviewing and Editing; RJNE: Conceptualization, Supervision, Funding acquisition, Writing - Reviewing and Editing. HNN and SK contributed equally.

## Abstract

Cytokinins (CKs) play a fundamental role in regulating dynamics of organ source/sink relationships during plant development, including flowering and seed formation stages. As a result, CKs are key drivers of seed yield. The cytokinin oxidase/dehydrogenase (CKX) is one of the critical enzymes responsible for regulating plant CK levels by causing their irreversible degradation. Variation of *CKX* gene activity is significantly correlated with seed yield in many crop species while in soybean (*Glycine max* L.), the possible associations between *CKX* gene family members (GFMs) and yield parameters have not yet been assessed. In this study, seventeen *GmCKX* GFMs were identified, and natural variations among *GmCKX* genes were probed among soybean cultivars with varying yield characteristics. The key *CKX* genes responsible for regulating CK content during seed filling stages of reproductive development were highlighted using comparative phylogenetics, gene expression analysis and CK metabolite profiling. Five of the seventeen identified *GmCKX* GFMs, showed natural variations in the form of single nucleotide polymorphisms (SNPs). The gene *GmCKX14*, with high expression during critical seed filling stages, was found to have a non-synonymous mutation (H105Q), on one of the active site residues, Histidine 105, previously reported to be essential for co-factor binding to maintain structural integrity of the enzyme. Soybean lines with this mutation had higher CK content and desired yield characteristics. The potential for marker-assisted selection based on the identified natural variation within *GmCKX14*, is discussed in the context of hormonal control that can result in higher soybean yield.

**Key Message:** Natural variations in soybean cytokinin oxidase/dehydrogenase gene, *GmCKX14*, with high expression during seed development, were linked to increased sink strength via altered cytokinin profiles in high yielding cultivars.

## Introduction

Soybean (*Glycine max* L.) is a legume species that is cultivated on a global scale, and is important to food and biomaterials industries (Liu et al. 2020). The high oil and protein content of mature seeds make it a valuable crop. However, soybean cultivation performs well below its full potential due to high rates of flower and pod abortion during reproductive development (Nico et al. 2016; He et al. 2019). Advanced genomics techniques including gene mapping, genome wide association studies, marker development and molecular breeding can be used to enhance the efficiency of classic selection and accelerate soybean yield improvement (Hua et al. 2019; Afzal et al. 2020). Thus, employing genomic studies for the identification and characterization of relevant genes and the development of molecular markers responsible for yield (e.g. higher rates of flower and pod set, increased number of seeds and 100 seed weight) is important for achieving a Green Revolution in soybean (Liu et al. 2020). Notably, previous research has reported a significant role for certain signaling molecules, the cytokinins (CKs), in regulating flower abortion rates and for increasing the number of pods being set in several crop species (Carlson et al. 1987; Atkins and Pigeaire 1993; Nonokawa et al. 2012; Emery et al. 2000; Emery and Atkins 2006). This includes soybean, as enhanced CK content was reported in reproductive tissues during pod set and seed filling phases (Jameson and Song 2016; Liu et al. 2018; Zuñiga-Mayo et al. 2018).

Cytokinins are adenine derivatives which are differentiated based on side chain substitutions at the N^6^ position of the purine ring. Four primary isoprenoid CKs occur in nature: isopentenyladenine (iP), *trans*-zeatin (*trans*Z), *cis*-zeatin (*cis*Z) and dihydrozeatin (DZ) (Sakakibara 2006). The occurrence and physiological activity of CKs can differ not only among plant species, but also among different plant organs (Gajdosová et al. 2011). CK homeostasis is balanced through the rate of their biosynthesis, interconversion, inactivation and degradation (Sakakibara 2006). The rate limiting step in CK biosynthesis is regulated by an enzyme isopentenyl transferase (IPT), while their breakdown is mainly controlled by another enzyme, cytokinin oxidase/dehydrogenase (CKX) (Hirose et al. 2008). CKX catalyzes CK degradation by cleavage of the N^6^-isoprenyl sidechain (Kieber and Schaller 2014).

The evolutionary development of the CK-catabolizing genes and their distinct individual properties indicate there are important roles for CKX enzymes in CK regulation (Wang et al. 2020). In plant species, CKX enzymes are produced by multi-gene families. Recently, progress has been made in identifying the mechanism of *CKX* gene family members (GFMs) to improve seed grain yield in several important crops, such as rice (*Oryza sativa* L.) (Gao et al. 2014; Gao et al. 2019; Mao et al. 2020), barley (*Hordeum vulgare* L.) (Zalewski et al. 2014; Holubová et al. 2018; Gasparis et al. 2019), and wheat (*Triticum aestivum* L.) (Zhang et al. 2012; Lu et al. 2015; Chang et al. 2015; Chen et al. 2020), making *CKX* an emerging genetic target for crop yield improvement (Jameson and Song 2016; Szala et al. 2020; Chen et al. 2020). Considering the potential of *CKX* genes, Le et al. (2012) identified and characterized expression of soybean *CKX*s at various plant development stages. However, no functional studies and no genetic markers for yield in relation to CKs have been reported in soybean to date.

An enhanced level of endogenous CKs in plants is often a result of loss of function or knock out of *CKX* genes, that can lead to increased seed numbers and seed weights (Chen et al. 2020). In Arabidopsis (*Arabidopsis thaliana* L.), *AtCKX3/AtCKX5* double mutants formed larger inflorescences, with increased flowers and seed numbers (Bartrina et al. 2011). In cotton (*Gossypium* spp.), moderate suppression of *CKX* genes led to higher yields of seed and fiber and this was brought about by several CK controlled traits including: delayed leaf senescence, increased photosynthetic activity, enhanced formation of fruiting branches and bolls, and enlarged seed size (Zhao et al. 2015). A similar effect has been documented in barley, for which silencing of *HvCKX1* and *HvCKX9* stimulated seed sink strength, and effectively increased grain weight (Zalewski et al. 2010, 2014). Subsequently, silencing of *HvCKX1* via RNAi led to enhanced plant productivity (Holubová et al. 2018).

Landmark reports in CK research using rice and wheat reported that natural genetic variation within *CKX* genes can significantly impact yield traits. One of the first reports on *CKX*-associated yield increases was a natural variant of rice *OsCKX2* gene which had a premature stop codon (Ashikari et al. 2005). Association of *OsCKX2* activity with grain number was further confirmed by knocking out its expression which resulted in increased tiller number and improved yield, as well as reduced yield loss under salinity stress conditions (Joshi et al. 2018). Similarly, Yeh et al. (2015) reported that reduced expression of *OsCKX2* resulted in higher CK accumulation in reproductive organs, thereby enhancing grain yield. Furthermore, the natural variation in *TaCKX6-D1* – a wheat ortholog of rice *OsCKX2*, was significantly associated with grain weight (Zhang et al. 2012). Novel allelic variation within another wheat *CKX* gene - *TaCKX6a02*, was also correlated with grain size, weight, and seed filling rate (Lu et al. 2015).

These functional studies suggest that, though plant *CKX* genes belong to multigene families, loss of function of a single gene that is expressed at critical stages of seed development can result in a dramatic yield change. Therefore, manipulation of CK metabolism by modifying the *CKX* activity can be effectively used to enhance yield in important crop species. The cultivation of genetically modified crops has become a contentious topic because of a range of issues related to food security, environmental risks and human health (e.g. gene dispersal, loss of biodiversity, the emergence of superweeds and superpests, raising antibiotic resistance, food allergies and other side effects; Gassmann et al. 2011; Ahmad and Mukhtar 2017). As these matters will be difficult to resolve in the foreseeable future, the identification and exploitation of natural variation resources becomes an increasingly valued approach for breeding programs with the purpose of improving crop yields, especially for cases when GMO technology is either not available or meets with consumer or industry adoptive reluctance. The analysis of the single nucleotide polymorphisms (SNPs) is one of the effective techniques of exploring natural genetic variation in plants. Each SNP represents a difference in a single DNA base which can effectively change the phenotype of the modified alleles if they occur at functionally important locations (Mammadov et al. 2012; Su et al. 2019).

This work reports on a multi-pronged study of *CKX* gene family members in soybean including phylogenetic analysis, screening for natural variations, transcript expression and CK abundances among cultivars. A total of sixteen soybean cultivars with varying yield parameters were probed for natural variations in *GmCKX* genes. A number of single nucleotide polymorphisms (SNPs) were identified and evidence for association between a non-synonymous variation in *GmCKX14* and soybean yield parameters is presented. The outcome is a natural variant marker of CK metabolism for immediate use in plant breeding programs or, for further genetic engineering to develop new, high-yielding soybean cultivars.

## Materials and Methods

### Plant materials and growth conditions

Sixteen soybean cultivars - 9 large size beans and 7 natto beans (Sevita International, Canada) with varying seed sizes and yield levels (**Table S1**) were used in this study. Soybean plants were grown in the Trent University Aurora research greenhouse (Conviron, Canada; photoperiod 16/8h, 27/19 °C, 60-80% relative humidity (RH)). Seeds were planted in Sunshine Mix #1, professional growing mixture (Sun Gro Horticulture, Canada). Plants were watered daily and fertilized with Jack’s 15-16-17 fertilizer (JR Peters, USA) twice per week until the initiation of reproductive development. Leaves, pods and/or seeds were collected at developmental stages specific to the experiments. Stages of seed development were determined by visual appearance following descriptions previously indicated [https://extension.umn.edu/growing-soybean/soybean-growth-stages] including: R4 (full pod) at 20-30 days after flowering, R5 (beginning seed) at 30-45 days after flowering, R5.5 (intermediate stage between R5 and R6), and R6 (full seed) at 45-65 days after flowering.

### Identification of *CKX* genes in soybean and phylogenetic analysis

In order to identify the soybean *CKX* GFMs, full length sequences of the previously identified genes of *Arabidopsis thaliana CKXs* and *Oryza sativa CKXs* (Gu et al. 2010) were used as a query to search the soybean genome database in phytozome v9.1 (http://phytozome.net/soybean.php) (Schmutz et al. 2010). The 17 putative soybean *CKX* genes were confirmed based on the presence of domain signatures (Schmülling et al. 2003), necessary for enzyme function, e.g. FAD binding domain (PF01565) and CK binding domain (PF09265) against the Protein Family (PFAM) database (Finn et al. 2014). Gene nomenclature was based on the chromosome on which they occurred and, in case of tandem duplications, the order of their location. The identified *GmCKX* GFMs along with their respective gene and protein characteristics are presented in **Table 1**.

**Table 1.** Characteristics of the analysed soybean *CKX* genes and their respective proteins.

| Gene name | Accession no. | No. of Exons | Size of gene (bp) | Size of AA sequence | Mw (KDa)<sup>a</sup> | pI<sup>a</sup> | Glycosylation sites<sup>b</sup> |
| --- | --- | --- | --- | --- | --- | --- | --- |
| *GmCKX03* | Glyma03g28910 | 7 | 2757 | 545 | 61.1 | 6.71 | 2 |
| *GmCKX04-1* | Glyma04g03130 | 5 | 6403 | 451 | 51 | 6.47 | 1 |
| *GmCKX04-2* | Glyma04g05840 | 6 | 3413 | 494 | 55.1 | 5.22 | 2 |
| *GmCKX06* | Glyma06g03180 | 5 | 7205 | 528 | 59.5 | 6 | 1 |
| *GmCKX09-1* | Glyma09g35950 | 6 | 2998 | 534 | 60.1 | 6.37 | 3 |
| *GmCKX09-2* | Glyma09g07360 | 6 | 4028 | 536 | 60.8 | 9.22 | 2 |
| *GmCKX09-3* | Glyma09g07190 | 6 | 4934 | 533 | 60 | 6.95 | 6 |
| *GmCKX11* | Glyma11g20860 | 5 | 2997 | 552 | 62.3 | 6.79 | 6 |
| *GmCKX12* | Glyma12g01390 | 5 | 2908 | 442 | 49.2 | 6.46 | 1 |
| *GmCKX13-1* | Glyma13g16430 | 5 | 4042 | 535 | 61 | 7.35 | 1 |
| *GmCKX13-2* | Glyma13g16420 | 5 | 4829 | 524 | 58.8 | 4.95 | 7 |
| *GmCKX14* | Glyma14g11280 | 4 | 5891 | 513 | 57.2 | 5.81 | 3 |
| *GmCKX15* | Glyma15g18560 | 6 | 4071 | 543 | 61.5 | 8.73 | 2 |
| *GmCKX17-1* | Glyma17g06220 | 5 | 4175 | 535 | 60.9 | 6.42 | 2 |
| *GmCKX17-2* | Glyma17g06230 | 6 | 4598 | 528 | 59.5 | 5.41 | 6 |
| *GmCKX17-3* | Glyma17g34330 | 4 | 7149 | 513 | 57.4 | 6.16 | 2 |
| *GmCKX19* | Glyma19g31620 | 5 | 2596 | 544 | 60.7 | 6.12 | 4 |

<sup>a</sup> Molecular Weight and Isoelectric Point of the CKX proteins were predicted using the pI/Mw tool in ExPASy (http://web.expasy.org/compute_pi/)
<sup>b</sup> Potential glycosylation sites were predicted using NetNGlyc (http://www.cbs.dtu.dk/services/NetNGlyc/)

For phylogenetic analysis, the soybean CKX protein sequences along with previously reported protein sequences of the CKX proteins from the model plant species Arabidopsis, and two crops, rice and corn (Gu et al. 2010) were first aligned using the ClustalW interface in MEGA software (Penn State University, USA) (gap open penalty of 10 and gap extension penalty of 0.2; Rédei 2008; Hall 2013). The neighbourhood-joining method adopted from Gu et al. (2010) was used to construct a phylogenetic tree. Pair-wise deletion was applied to circumvent the gaps and Poisson correction was used to estimate the evolutionary distance. A bootstrap test with 1000 bootstrap replicates was conducted to check the reliability of the clusters.

### DNA isolation and amplification of *GmCKX* genes

Young leaves were collected from each soybean cultivar at the third trifoliate stage (V3). Approximately, 100-150 ng/μL of DNA was extracted from 100 mg of ground leaf tissue using DNeasy Plant Mini Kit (Qiagen Inc., Canada) following the manufacturer’s protocol. The levels of DNA in the extracted samples were quantified using NanoDrop 8000 (Fisher Scientific, Canada) spectrophotometer. All extracted DNA samples were diluted to a standard concentration of 30 ng/μL prior to further analysis.

Degenerate primers specific to each gene were designed using Primer3plus software [https://primer3plus.com/] for all 17 identified *CKX* genes following the guidelines for primer design provided by the PCR amplification kit manufacturer (Promega Corporation, USA; **Table S2**). PCR amplification of the identified *CKX* GFMs was performed using the reaction kits selected based on the gene size. Genes smaller than 4kb were amplified using the GoTaq^®^ Flexi DNA polymerase kit (Promega Corporation, USA). Cycling conditions were: 95 °C for 1 min, 35 cycles (95 °C for 10 s, 50-59 °C for 5 s and 72 °C for 2-4 min), 72 °C for 5 min, followed by a hold at 4 °C. Genes bigger than 4kb were amplified using Expand Long Range dNTPack (Roche, Switzerland). Cycling conditions were: 92 °C for 2 min, 10 cycles (92 °C for 10 s, 50-57 °C for 5 s and 68 °C for 2-7 min), 25 cycles (92 °C for 10 s, 50-55 °C for 5 s and 68 °C for 4.5-7 min), 68 °C for 7 min, followed by a hold at 8 °C. Detailed information on the annealing temperature and extension time for each of the 17 genes is included in **Table S2**. PCR products were purified using the QIAquick PCR purification kit (Qiagen Inc., Canada), quantified using a NanoDrop™ 8000 (Fisher Scientific, Canada) spectrophotometer, and diluted to a standard concentration of 20ng/μL.

### Sequencing analysis of *GmCKX* genes and identification of single nucleotide polymorphisms (SNP)

Sequencing primers were designed for every 400 bp, excluding larger intron regions, with Primer3plus software [https://primer3plus.com/] (**Table S3**). To amplify the 400bp regions, sequencing reaction mixes were prepared using the Big Dye Terminator V3.1 Cycle Sequencing Kit (Fisher Scientific, Canada). Cycling conditions were: 96 °C for 1 min followed by 35 cycles of 96 °C for 10 s, 50 °C for 5 s and 60 °C for 4 min. To determine if the PCR products were derived from a single locus or from two or more homoeologous regions, the resultant products were separated using a 3730 DNA Analyzer (Fisher Scientific, Canada) and analysed using Sequence Analysis software V5.4 (Fisher Scientific, Canada).

The sequences obtained from 16 soybean cultivars for each of the 17 *CKX* genes were compared to the reference genome from the phytozome v9.1 database using the MEGA software (Penn State University, USA) for any differences. The electropherograms of the identified differences were manually verified to eliminate ambiguity in nucleotide base calls. The regions of single nucleotide polymorphism (SNP) from the cultivars that showed differences from the reference gene sequence were re-sequenced along with 2 control cultivars, using fresh plant material and newly extracted DNA, in both forward and reverse direction with new primers (**Table S4**) to confirm the presence of SNPs.

### Predicted functional effects of the identified SNPs

To examine the potential protein structure of SNPs, swiss-models were constructed using Expert Protein Analysis System (ExPASy; Swiss Institute of Bioinformatics, Switzerland; Biasini et al. 2014). A PDB file of the constructed protein model was analysed using PyMol v2.0 software (The PyMOL Molecular Graphics System 2.0, Schrodinger, LLC., https://pymol.org/2/). The SNP location was identified in the primary protein structure and any effects were predicted based on the properties of the substituted amino acid (Betts and Russell 2007). Protein stability was predicted using the Site Directed Mutator software (SDM; UK; Topham et al. 1997; Pandurangan et al. 2017).

### Expression analysis of *GmCKX* genes by real-time qPCR

Total RNA was extracted from soybean seeds (OAC Wallace) at 3 developmental stages: R5, R5.5, and R6, using RNeasy Lipid Tissue Mini Kit (Qiagen Inc., Canada) and treated with DNase I (Qiagen Inc., Canada) before reverse transcription and cDNA synthesis to remove any genomic DNA contamination (Song et al. 2012). RNA samples were quantified using Nanodrop 8000 (Fisher Scientific, Canada) before reverse transcription. The first strand cDNA was synthesized using Omniscript Reverse Transcriptase Kit (Qiagen Inc., Canada) and Oligo-dT primers (Integrated DNA Technologies, Inc., Canada). The first strand cDNA and the gene-specific primers of 11 *GmCKX* genes were used in qRT-PCR reactions (**Table 5**). Primers used for qRT-PCR were designed with Eurofins’ Primer Design Tools (Eurofins, North York, ON, Canada) using following parameters: primer sequence length: 20-22 nt, GC content: 50% and 55%, melting temperature (Tm): 55 – 60 °C, and the size of the amplicon: 90-100 bp. The GoTaq qPCR Master Mix Kit (Promega Corporation, USA) was used for the gene expression profiling. The housekeeping gene *GmGAPDH* was used as an endogenous control for normalization of expression analysis. The qRT-PCR was performed using a StepOne Plus Real-Time PCR Thermo-cycler (Fisher Scientific, Canada), with an initial GoTaq^®^ Hot Start Polymerase activation step at 95 °C for 2 minutes, followed by 40 cycles of denaturation at 95 °C for 15 seconds, then annealing and extension at 60 °C for 1 minute under the Standard Cycling Conditions.

The expression levels of the *GmCKX* genes were tested in 3 biological replicates and each reaction was analysed in 2 technical replicates. Relative gene expression levels and fold changes were determined by comparison with the most highly expressed *GmCKX* gene for each corresponding seed developmental stage (Le et al. 2012). The differences at each growth stage and between the stages were compared statistically for significance (ANOVA and Duncan’s multiple range test; *p*<0.05).

### Cytokinin extraction and quantification using High Performance Liquid Chromatography – Electrospray Ionization - Tandem Mass Spectrometry (HPLC-(ESI+)-MS/MS)

Cytokinin profiles were analysed from pods (collected at R4 and R5 stage) and seeds (collected at R5.5 and R6 stages) of 4 soybean cultivars: OAC 06-34, DH420, OAC Wallace and DH3604. Each sample type was analysed in 3 biological replicates. The CK extraction procedure was performed as previously described (Farrow and Emery 2012). Profiles of 25 CKs were scanned for and analyzed, if present, including: four CK free bases (FB), their corresponding riboside (RB), nucleotide precursors (NT), nine glucoside (GLUC) and four 2-methylthiol conjugates (MET).

Purified CKs were identified and quantified by HPLC-(ESI+)-MS/MS (Agilent 1100 series HPLC (Agilent, USA) connected to a QTrap 5500 mass spectrometer (Sciex, USA)), according to the conditions outlined in Farrow and Emery (2012). An aliquot was injected on a Luna C18 reversed-phase column (3 μm, 150 × 2.0 mm; Phenomenex, USA), and the CK were eluted with an increasing gradient of 0.08% CH_3_COOH in CH_3_CN (A) mixed with 0.08% CH_3_COOH in ddH_2_O (B), at a flow rate of 0.2 mL/min. The initial conditions were 5% A and 95% B, changing linearly in 17 min to 95% A and 5% B. Conditions remained constant for 5 min, and then, immediately returned to initial conditions for 18 min. The effluent was introduced into the electrospray source using conditions specific for each CK, and quantification was obtained by multiple reaction monitoring of the protonated intact CK molecule [M+H]^+^ and the specific product ion.

The differences in the levels of the select CK groups (FB, RB, NT) and in CK types (tZ, cZ, DZ and iP-type) belonging to these three groups were compared statistically for significance (ANOVA and Duncan’s multiple range test; *p*<0.05) among the 4 cultivars at each growth stage.

## Results

### Identification and genetic analysis of the *CKX* GFMs in soybean

The BLAST search report using Arabidopsis and rice *CKX* genes produced 17 putative soybean *CKX* genes (*GmCKX*s). Sequence analysis using the Protein Family PFAM database confirmed the presence of domain signatures required for enzyme function (i.e. FAD binding and CK binding domains) in all 17 soybean *CKX* homologs.

The identified soybean *CKX* genes were predicted to be located on 10 different chromosomes and disrupted by introns. All *GmCKX* genes displayed a minimum of 4 exons, with a few having 6 and 7 exons, revealing a high complexity of the sequences (**Table 1**). The predicted *GmCKX* genes ranged from 2596 to 7205 bp, with amino acids ranging from 442 to 552 aa per protein sequence, molecular weight ranging from 51.0 to 62.3 KDa, and theoretical isoelectric point ranging from 4.95 to 9.22. The predicted molecular weights and isoelectric points of their proteins are similar to those of other plant species exhibiting similar protein functions. Soybean *CKX* GFMs were predicted to have 1 to 6 glycosylation sites (**Table 1**) suggesting post translational modification and a broad range of functions in plants.

### Phylogenetic analysis of *GmCKX* GFMs

To dissect the phylogenetic relationships of the *GmCKX* GFMs, a multiple alignment analysis was performed using the full amino acid sequences of the *CKX* GFMs of soybean, Arabidopsis, rice and corn. Based on the alignment analysis, *GmCKX* genes were distributed into 4 out of 6 groups of the generated phylogenetic tree – I, III, IV and V (**Fig. 1**, **Fig. S1)**with well supported bootstrap values. Group I consisted only of dicot genes, groups II and VI consisted of only monocot genes, while groups III, IV and V contained both dicot and monocot genes. This is congruent with a similar set of 6 groups that was generated in a phylogenetic study on corn *CKX* GFMs (Gu et al. 2010)

**Fig. 1.**
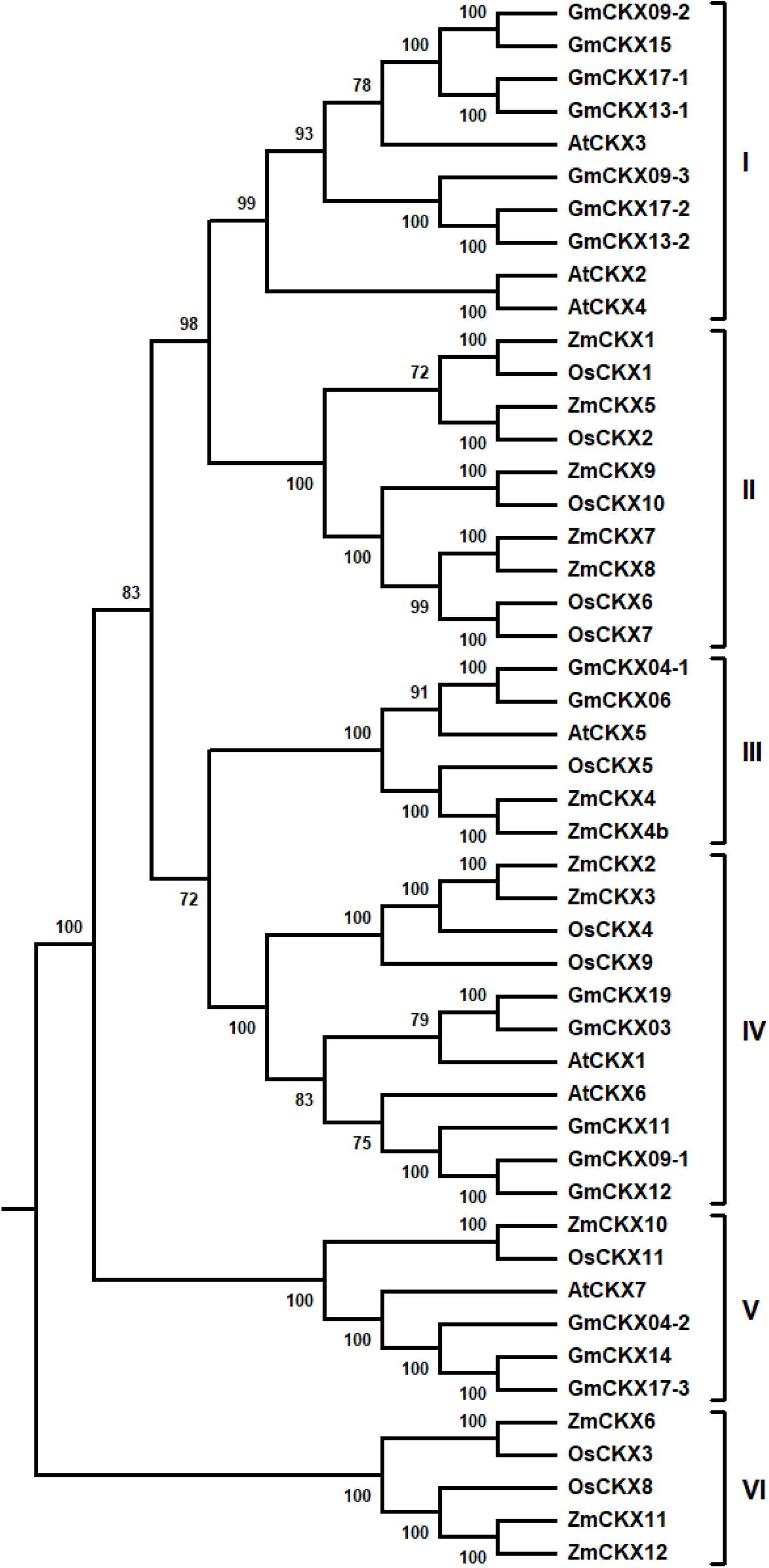
Phylogenetic analysis of the CKX gene family members of two monocot species, rice (*Oryza sativa*), maize (*Zea mays*), and two dicots, Arabidopsis (*Arabidopsis thaliana*) and soybean (*Glycine max*). The tree was generated based on the protein sequences, upon alignment with ClustalW, using the neighbor-joining method. The CKX proteins were assigned into 6 groups of which four (I, III, IV and V) contained GmCKX proteins and the other two (II and VI) were exclusive to monocots

### *Gm*CKX gene sequencing and identification of single nucleotide polymorphism (SNP)

To identify potential variations in *GmCKX* GFMs, complete sequencing of 17 *GmCKX* genes was performed from 16 soybean cultivars. Among the natural variation detected in the analysed genes, total of 9 non-synonymous SNPs were found in 5 *GmCKX* genes (*GmCKX04-1, GmCKX04-2, GmCKX06, GmCKX09-1, GmCKX14*). Resultant amino acid changes of the identified SNPs are listed in **Table 2** along with their potential destabilizing effect (as predicted by SDM, see methods for details) and the cultivars in which each SNP is present.

**Table 2.** *GmCKX* gene family members with natural variation and the cultivars in which the variations occurred. Information associated with SNPs found in *GmCKX14* is bolded (more description in text).

| Gene | Variation | Pseudo $\Delta\Delta G^a$ | Cultivar |
| --- | --- | --- | --- |
| *GmCKX04-1* | T419P | 0.60 | DH3604, X5076 108B, OAC 06-14 |
| *GmCKX04-2* | A66V | -1.57 | DH3604, X5076-108B |
| *GmCKX06* | H16R<br>K77E | -0.63 | X5076 68B, X5076 159B |
| *GmCKX09-1* | L330F<br>L509Q | -0.34 | DH3604, OT 08 13, X5076 159B |
| *GmCKX14* | **T69A**<br>**H105Q**<br>**A159G** | **-0.75** | **OAC Wallace, IA1010LF,**<br>**DH420, DH748** |

<sup>a</sup> $\Delta\Delta G$ value as per SDM classifies a mutation as slightly destabilizing if less than -0.5; slightly stabilizing between 0.5-1.0; and destabilizing if less than -1.0

The *GmCKX14* sequence showed 8 nucleotide variations within the first exon, present in 4 regular bean type cultivars: OAC Wallace, IA1010LF, and DH420 - characterized by high seed weight (>20 g per 100 seeds), and in high yielding DH748 (over 4 t/ha; **Table S1**). Out of the 8 SNPs identified in *GmCKX14*, 3 were non-synonymous and resulted in amino acid changes: T59A, H105Q and A159G (**Table 2**). A comparison of the homology model proteins of the two haplotypes created by ExPASy and a prediction of the effect of variation on the protein using SDM revealed that the combined amino acid changes affects the stability of the enzyme produced by the modified *GmCKX14* gene (ΔΔG of −0.75). Moreover, one of these variations (H105Q), is within an active site residue, histidine, that was previously reported (Kopečný et al. 2015) to be responsible for the co-factor (Flavin Adenine Nucleotide, FAD) binding and maintaining the enzyme’s structural integrity. This substitution could also affect the activity of the CKX enzyme by reducing its affinity to the CK substrate, similar to prior *in vitro* studies where a maize CKX variant H105A resulted in 100 fold lower activity compared to wildtype (Kopečný et al. 2015), hence leading to higher CK accumulation and subsequently improved yield parameters.

Based on the presence of the destabilizing SNPs, 2 cultivars, OAC Wallace and DH420 were selected for further analysis of CK metabolites and compared to 2 other cultivars that did not have the *GmCKX14* SNP. In addition, one cultivar, OAC Wallace was used for the expression analysis of the selected *GmCKX* GFMs.

### Expression patterns of Gm*CKX* GFMs during soybean reproductive development

In the developing soybean seeds of OAC Wallace, *CKX* GFMs, 6 of the eleven analysed *GmCKX* genes had relatively high expression levels at R5, R5.5, and R6 growth stages: *GmCKX14, GmCKX03, GmCKX19, GmCKX04-1, GmCKX04-2*, and *GmCKX15* (**Fig. 2**). *GmCKX14* had, by far, the highest expression level in all three seed stages compared to the other *GmCKX*s (ANOVA, Duncan’s multiple range test, *p*<0.05). Additionally, the transcripts of *GmCKX14* and *GmCKX19* showed clear, increasing trends along the seed development progression from R5 to R6 stage (ANOVA, Duncan’s multiple range test, *p*<0.05).

**Fig. 2.**
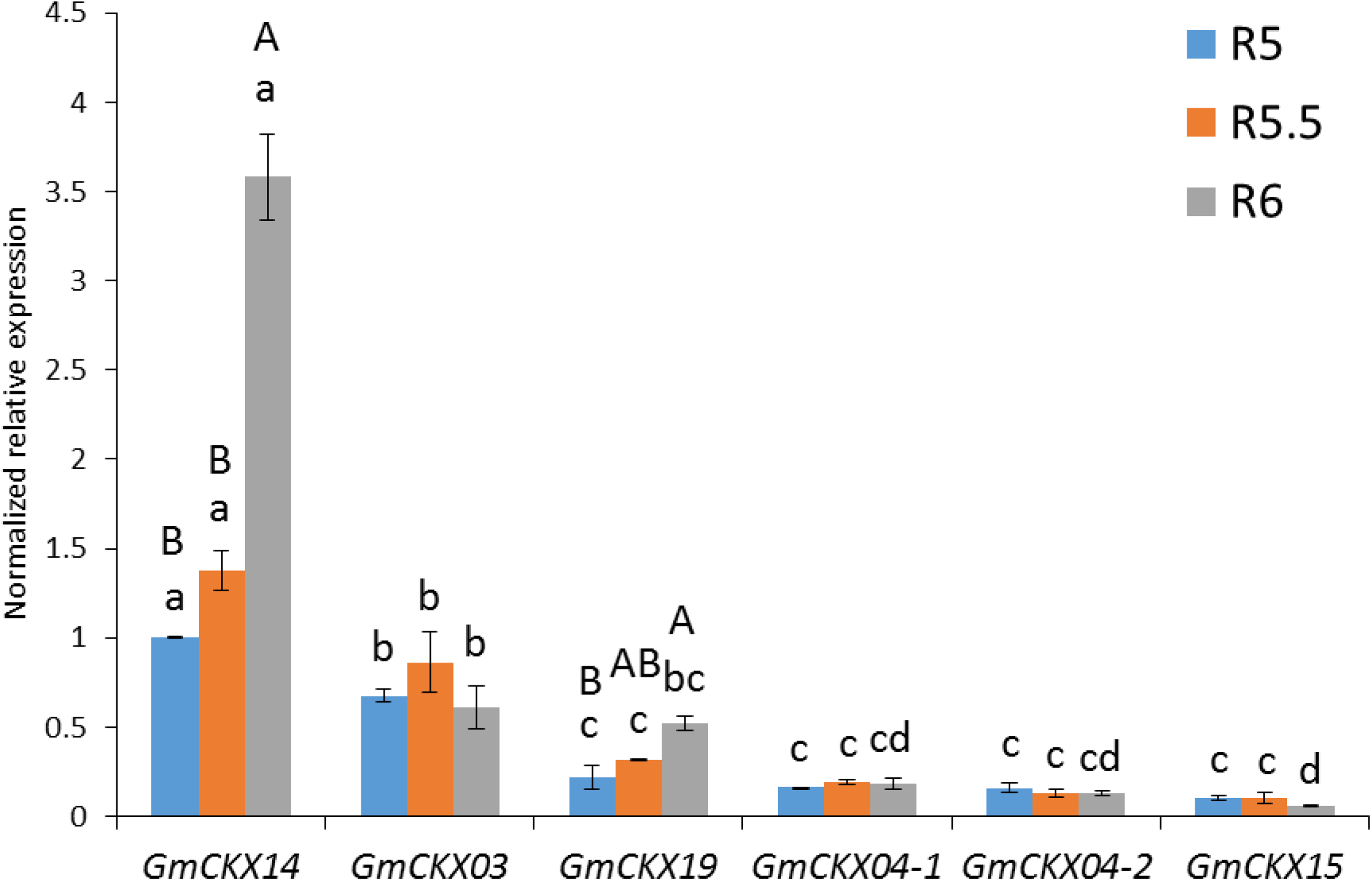
Quantitative expression profiles of 6 *GmCKX* genes analysed at 1 soybean developmental stages: R4 (pods), R5.5 (seeds) and R6 (seeds) in OAC Wallace. The housekeeping gene *GmGAPDH* was used as an endogenous control for normalization of expression analysis. The relative transcript level is presented using 2 delta CT method (qRT-PCR). Error bars represent standard error of the mean (n=3). Small letters denote significant differences among the genes at each of the growth stages and capital letters denote significant differences among the three stages for particular gene (ANOVA, Duncan’s multiple range test, *p*<0.05)

### Cytokinin quantification by HPLC-MS/MS during soybean reproductive development

The levels of 25 forms of CKs were measured during soybean seed development using HPLC-MS/MS (**Fig. 3**, **Table S6a-b**). Notably, distinct patterns in the CK form and levels were recorded across the 4 stages of pods (R4, R5) and seeds (R5.5, R6) among the 2 high yielding cultivars, OAC Wallace and DH420 (both of which have the *GmCKX14* SNPs), and the other 2 analysed cultivars (OAC 06-14 and DH3604 – which do not have the *GmCKX14* SNPs). The observed trends are consistent with the reduced *GmCKX14* activity. For example, DH420 had the highest level of CK precursors, NTs, in pods at R4 and R5 stages (ANOVA, Duncan’s multiple range test, *p*<0.05). In OAC Wallace, at the later R6 stage, the FB CK levels were the highest among the tested soybeans (ANOVA, Duncan’s multiple range test, *p*<0.05). This strongly aligns with the results of gene expression analysis for *GmCKX14* in R6 seeds of this cultivar, since *GmCKX14* at R6 was the most highly expressed gene of any timepoint. Regarding CK-type (cZ, tZ, DZ and iP-type) distribution, the highest yielding cultivar, OAC Wallace, revealed the highest accumulation of the most active, tZ-type CKs in seeds at both R5.5 and R6 stages (ANOVA, Duncan’s multiple range test, *p*<0.05). Additionally, at R6 stage, cZ levels in OAC Wallace exceeded those in the remaining 3 cultivars (ANOVA, Duncan’s multiple range test, *p*<0.05). Cytokinin-type profiles were different for the second *GmCKX14* SNP cultivar, DH420, but there were still remarkable patterns which may explain its high yield. Specifically, the levels of cZ and iP derivatives were the most abundant at R4 compared to the other 3 soybeans.

**Fig. 3.**
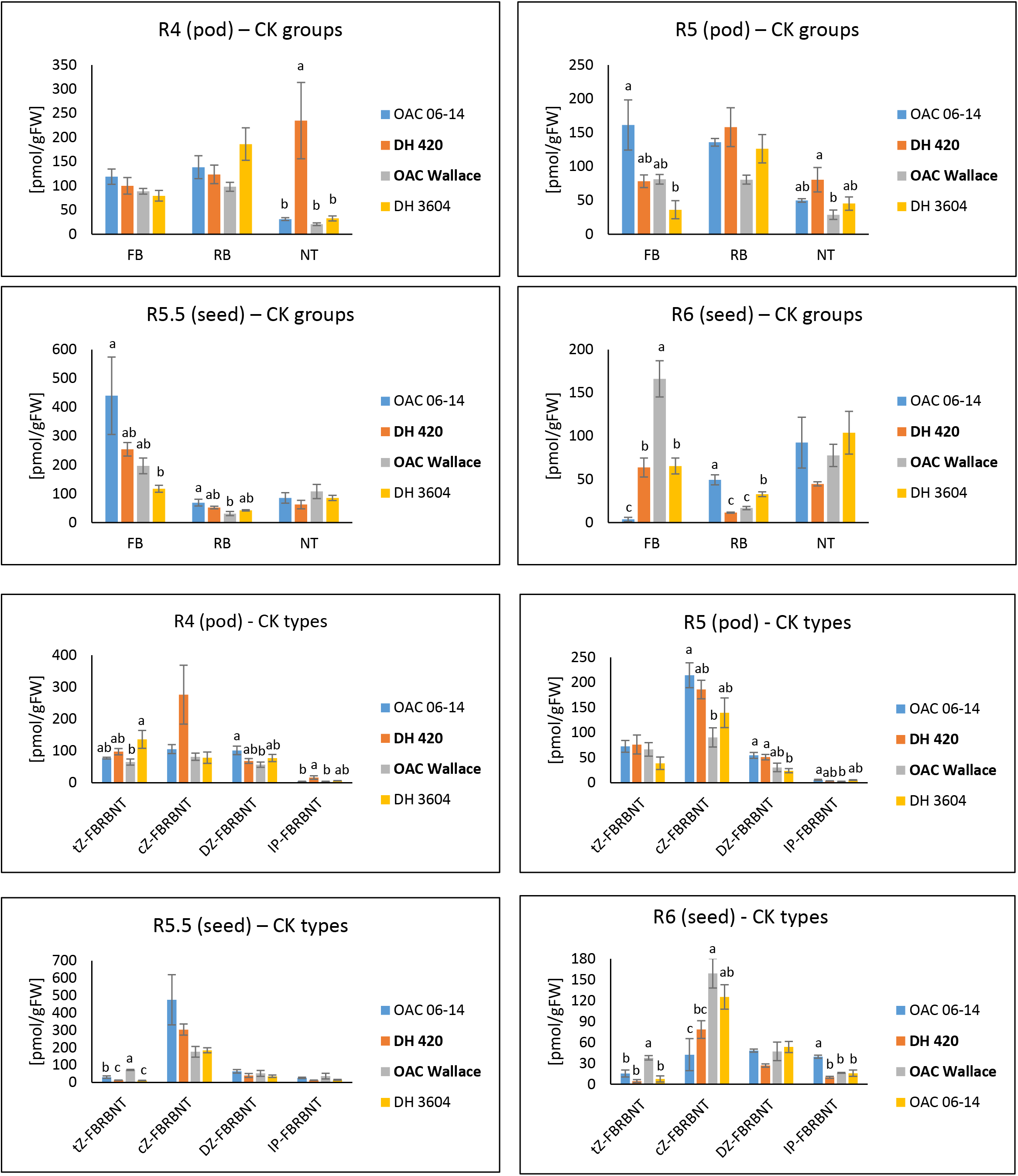
CK profile data (groups: FB, RB and NT, and types within these groups: tZ-type, cZ-type, DZ-type and iP-type) analysed using HPLC-MS/MS at 4 developmental stages in 4 soybean cultivars. Highlighted bold are the cultivars with SNPs in *GMCKX14* detected. Letters denote significant differences among the cultivars (ANOVA, Duncan’s multiple range test, *p*<0.05)

## Discussion

Cytokinins (CKs) play essential roles in numerous plant physiology processes. Among their multiple functions, CKs are a key drivers of seed yield (reviewed by Jameson and Song 2016; Chen et al. 2020). In soybean, CKs are: involved in preventing flower abortion and in pod setting (Nonokawa et al. 2012); associated with seed development and fatty acid biosynthesis (Nguyen et al. 2016); and ultimately, responsible for seed yield (Kambhampati et al. 2017). Studies on genetic analysis of CKs aid greatly in the understanding of fundamentals of plant physiology and they help build genetic resources for high yielding crop breeding programs. Although CK metabolic genes have been extensively studied in several plant species (Chen et al. 2020; Wang et al. 2020a, b), only a few examples have investigated the function of CK metabolic genes in soybean yield formation (Le et al. 2012). Thus, the genetic characterization and identification of CK metabolic genes, along with the selection of unique natural variants for high-yielding traits should be a high priority for the progress in soybean breeding. To date, the present study of *GmCKX* genes is the most focused example of the drive towards the identification of CK-based markers for modern soybean breeding programs.

This work focuses on the soybean CKX GFMs because it is the best-known way to effectively regulate endogenous CKs in reproductive tissues, and thus improve yield. While IPT is the limiting step for CK biosynthesis, it is challenging to control, and no variants that are linked to high yield were identified in crop species. By contrast CKX can degrade CKs in a spatially controlled manner and examples of both natural variants (e.g. Ashikari et al. 2005) and manipulation through genetic engineering (e.g. Zalewski et al. 2014) for enhanced crop yields are convincing. Our work employed a multi-pronged approach to study CK metabolism and its mechanistic association with soybean yield parameters. This included a *CKX* protein sequence based phylogenetic analysis, SNP identification in *CKX* GFMs among different yielding cultivars, bioinformatic predictions/literature comparison of SNP destabilization potentials, and *CKX* gene expression analyses as well as comprehensive CK metabolite profiling during pod and seed filling.

In this study, 17 *CKX* GFMs in soybean were identified (**Fig. S1**). Previously, the presence of paralogous *CKX* genes were reported from multiple species including rice (11 *CKX* GFMs; Mameaux et al. 2012), foxtail millet (11 *CKX* GFMs; Wang et al. 2014), corn (13 *CKX* GFMs; Zalabák et al. 2014), potato (5 *CKX* GFMs; Suttle et al. 2014), wild strawberry (8 *CKX* GFMs; Jiang et al. 2016), apple (12 *CKX* GFMs; Tan et al. 2018) and wheat (11 *CKX* GFMs; Chen et al. 2020), all of which were much lower numbers than those identified here for soybean. A higher number of *CKX* metabolic genes in the soybean genome compared to that of other species is likely a consequence of genome evolution and duplication, which produces homologous genes, increasing the total gene number (Kaltenegger et al. 2018; Wang et al. 2020a).

A multiple alignment analysis of the *CKX* GFMs of rice, corn, Arabidopsis and soybean revealed 6 distinctive clusters that grouped the CK degradation genes (**Fig. 1**). Group I consisted of only dicot genes, and groups II and VI of only monocot genes, suggesting that a relative genetic divergence may have occurred in *CKX* genes during evolution and speciation of the two plant groups (Tan et al. 2018). Groups III, IV and V contained both dicot and monocot genes, indicating that the expansion of these genes occurred before the monocot/dicot divergence (Chaw et al. 2004; Hertweck et al. 2015).

A phylogenetic tree further revealed that most of the soybean *CKX* genes have counterparts in Arabidopsis. For example, 4 of the *GmCKX* GFMs: *GmCKX09-2*, *GmCKX13-1*, *GmCKX15*, *GmCKX17-1* and Arabidopsis *AtCKX3*; *GmCKX09-1, GmCKX11, GmCKX12* and *AtCKX6*; *GmCKX03, GmCKX19* and *AtCKX1*; *GmCKX04-2*, *GmCKX14*, *GmCKX17-3* and *AtCKX7*; *GmCKX04-1*, *GmCKX06* and *AtCKX5* belong to the same clades (**Fig. 1**), suggesting soybean and Arabidopsis *CKX* genes descended from the same ancestor. The fact that not every soybean *CKX* gene has a counterpart in Arabidopsis may reflect that these two species have undergone differential expansions.

The genes assigned to group V (*GmCKX04-2, GmCKX14, GmCKX17-3 and ZmCKX10, AtCKX7, OsCKX11*) were previously found to have essential roles in plant growth and development. A cytosolic corn enzyme, *ZmCKX10*, preferentially degrades cisZeatin (cZ) and has an abundant transcript in embryos, leaves and pedicels (Šmehilová et al. 2009). In Arabidopsis, *AtCKX7* displays high affinity towards cZ (Gajdošová et al. 2011) while the gain-of-function mutant of *AtCKX7* exhibited an increased CKX enzymatic activity, leading to CK deficiency and altered vascular development (Köllmer et al. 2014). *OsCKX11* coordinates source and sink relationship, thus regulating leaf senescence and grain number in rice (Zhang et al. 2020). In soybean, *GmCKX04-2* and *GmCKX14* were highly expressed in the leaves, roots and root hairs at reproductive stage under normal and drought conditions (Le et al. 2012). These reports suggest the *CKX* genes of phylogeny group V have critical effects on vegetative and embryonic growth, somatic embryogenesis, seed development, lateral root formation, petal cell identity, floral meristem identity and transition to flowering. Therefore, the polymorphisms among the soybean *CKX* genes would be most meaningful if found among the genes of group V (*GmCKX04-2*, *GmCKX14*, *GmCKX17-3*) because of their reproductive tissue specific expression, which would likely most impact soybean yield and seed weight.

Single nucleotide polymorphisms (SNPs) have attracted interest in molecular breeding because of their genome-wide availability and amenability for high throughput, cost-effective platforms (Rasheed et al. 2017, Su et al. 2019). Several reports by commercial representatives such as Monsanto (Eathington et al. 2007; Rosso et al. 2011), Syngenta (Ribaut and Ragot 2007), Dow AgroSciences (Ren et al. 2008; Mammadov et al. 2012) and Hi-bred (Zheng et al. 2008) have shown that development of SNPs as markers has enormous utility in plant breeding. The use of SNP markers has also been recognized for convenience due to their relative frequency and heritability among plant genotypes. Previous reports described the potential for SNP markers for practical use in soybean breeding (Song et al. 2013; Lee et al. 2015). For example, mutations in soybean D-myo-inositol 3-phosphate synthase 1 gene (*MIPS1*) lead to the increase in inorganic phosphorous and sucrose levels, and reduced phytate and raffinosaccharide accumulation in seeds, improving the nutritional value of the food products (Maupin et al. 2011).

To identify SNPs within the soybean *CKX* GFMs, sequencing of 17 *GmCKX* genes was performed from 16 soybean cultivars. A total of 9 non-synonymous SNPs were found among 5 *GmCKX* genes: *GmCKX04-1, GmCKX04-2, GmCKX06, GmCKX09-1* and *GmCKX14* (**Table 2**). We had hypothesized that the presence of SNPs in *GmCKX14* sequences could be critical for determining reproductive organ sink strength because *GmCKX14* is highly expressed during seed development stages (Le et al. 2012). It was predicted that the disruptive SNPs would be present in high seed weight (100 seed weight > 20g) soybean cultivars, like OAC Wallace, DH420 and IA1010LF. Homology modelling and previous reports on active site residues of CKX enzymes (Kopečný et al. 2015), support the hypothesis that reducing the GmCKX14 enzymatic activity can lead to the increased accumulation of CKs, resulting in the increased sink strength of the developing seeds. Phylogeny analysis suggested that *GmCKX14* might be involved in degradation of cZ-type CKs, similarly to its *AtCKX7* ortholog (Köllmer et al. 2014; Schäfer et al. 2015). The potential substrate specificity of the *GmCKX14* gene towards cZ and any lower activity caused by the observed SNP substitutions (**Table 2**) could lead to the higher cZ-type CK levels and hence, higher seed weight (Kudo et al. 2012; Powell et al. 2013; Hluska et al. 2016; Kambhampati et al. 2017). Therefore, a possible explanation for the higher seed weight observed in OAC Wallace, DH420 and IA1010LF cultivars, could be the potentially lower activity of the *CKX* enzyme caused by the substitutions in *GmCKX14* sequence: T69A, H105Q and A159G, which, according to our models, affect protein stability. In particular, one of these substitutions, H105Q, occurred within the protein active residue responsible for co-factor binding, likely rendering the protein unstable, as previously described (Kopečný et al. 2015). Previous reports of CK metabolite profiles further support this hypothesis, as they showed a positive correlation between soybean seed weight and levels of cZ (Kambhampati et al. 2017).

The manipulation of *CKX* GFMs can alter endogenous active CK levels, and substantially modify plant physiology and development (Jameson and Song 2016; Chen et al. 2020). Genetically modified plants, with CK profiles altered via targeting *CKX* GFMs, show enhanced tolerance to environmental stresses and improved yield, as it has been observed in: barley (Zalewski et al. 2014; Holubová et al. 2018), rice (Ashikari et al. 2005; Yeh et al. 2015; Li et al. 2016; Joshi et al. 2018) and wheat (Zhang et al. 2012; Li et al. 2018; Wang et al. 2018). In the present study, one of the cultivars, OAC Wallace, was selected for gene expression analysis since it was high yielding and it contained the SNPs in *GmCKX14* sequence. Accordingly, transcript analysis showed that that patterns of *GmCKX*s expression in developing soybean seeds differed among three reproductive growth stages (R5, R5.5, and R6; **Fig. 2**). *GmCKX14*, was exceptional since it had the highest transcript levels in each of the seed growth stages. Therefore, it can be considered as a high priority gene candidate for the further studies of the functional characteristics (CRISPR/Cas9 or overexpression) and as a target in modern breeding programs, particularly in regards to high yield and abiotic stress tolerance (Pospíšilová et al. 2016; Holubová et al. 2018; Gasparis et al. 2019). Our results align with another study that showed that *GmCKX14* had high expression in flowers, during early pod development and seed formation (Le et al. 2012). Under normal conditions, high expression of *CKX* genes reduces quantities of the active forms of endogenous CKs, and their transport (RBs) and conjugate (GLUCs) forms (Avalbaev et al. 2012). This results in the reduced organ/tissue sink strength, limited photosynthetic activity, altered distribution of carbohydrates and glycolytic enzymes and, ultimately, reduced seed growth (Werner et al. 2008; Prerostova et al. 2018). Therefore, in our work, the SNPs identified in the 4 high-yield soybean cultivars can conceivably diminish the activity of *GmCKX14*, and thereby enhance CK accumulation during the reproductive stages. This, would thus delay senescence, enhance photosynthesis and primordia formation, which, in turn, can increase the spike number, flower number and size, pod number, seed number and seed weight (Jameson and Song 2016; He et al. 2018; Panda et al. 2018; Szala et al. 2020). Taken together, our data support the hypothesis that the substitution in *GmCKX14* is closely linked to seed yield in soybean. These findings highlight that the natural variations observed in *GmCKX14* could be important markers for developing high-yielding cultivars in soybean breeding programs.

CK metabolites are strongly associated with the enhancement of soybean reproductive development. HPLC□MS/MS based CK quantifications have clearly established their distinct correspondence with cell division, cell expansion, and dehydration during embryo development, seed protein and lipid accumulation, and in the DNA endoduplication (Nguyen et al. 2016; Kambhampati et al. 2017). Therefore, the present study surveyed the complete set of CKs during three reproductive stages that critically define pod and seed development and yield formation in soybean. This was done for two high yield cultivars (OAC Wallace and DH420) which contained the *GmCKX14* SNPs and two lower-yielding cultivars (OAC 06-14 and DH3604) that did not (**Fig. 3**).

Distinct CK metabolite profile attributes were observed for the two high yield cultivars. Notably, in the cultivar DH420 (*GmCKX14* SNPs identified), the inactive CK precursors, NTs, were a dominant form in R4 and R5 stages. Furthermore, the peak in cZ and iP-type CKs observed in DH420 during early pod development suggest that potential activity of the altered *GmCKX14* is important for the control of plant development and grain yield, as was the case for a previously reported mechanism of *HvCKX1* action which modified barley productivity (Holubová et al. 2018). In another high yield cultivar, OAC Wallace, (also with the *GmCKX14* SNPs identified), the active FB forms significantly peaked at the R6 stage. Two CK groups (NTs and RBs), that are transportable and are precursors to the most active FB forms were observed to preferentially accumulate in developing pods, thereby representing sources for the higher FB content in seed (Witte and Herde 2020). The FB peak was previously reported in soybean seeds at R6, at the onset of mitotic cell division events (Lin et al. 2017). The activity of FBs strongly contributes to the sink strength and accumulation of assimilates in soybean seed development (Kambhampati et al. 2017). As predicted by the protein model, the *GmCKX14* SNPs can bring about an increased accumulation of the active CK types (tZ, cZ) during R5.5 and R6 stages in OAC Wallace (**Fig. 3**). High levels of these CKs accumulate in different plant organs at the reproductive stages (Le et al. 2012) and correlate with grain yield in soybean (Kambhampati et al. 2017). This indicates potential effect of *GmCKX* SNPs on CK metabolism and increased soybean productivity, and reinforces the idea that the SNPs identified in *GmCKX14* can serve as a new, highly effective marker in soybean breeding programs.

Our comprehensive CK metabolite analysis is one of the rare examples in crops, where detailed endogenous hormone profiling has been shown to be functionally important in relation to the natural variation of *CKX* genes. However, not all the data point clearly toward high yield traits. Some of the discrepancies found in this study among the metabolic, gene expression data and bioinformatics prediction of *GmCKX14* function can be at least partially explained by the redundancy effects of 17 *CKX* GFMs in soybean (Liu et al. 2008; Lakhssassi et al. 2017). There is also the challenge of profiling metabolites that are derived from whole tissues for which resolution cannot be discerned at sub-cellular levels. It is now known that CKs are precisely sensed at multiple sites at the plasma-membrane and endoplasmic reticulum (ER) (Kubiasová et al. 2020) and that CKX isoforms are likewise specifically located, most frequently on the ER (Niemann et al. 2018). Thus, different CKX proteins would control different cytokinin pools depending on their subcellular localizations, which our methods cannot resolve.

Finding and exploiting genomic DNA variation is of utmost importance for crop genetics and breeding. In this research, soybean *CKX* gene activity has been demonstrated to be significantly associated with yield components. Gene sequencing in multiple soybean cultivars and bioinformatics predictions revealed that SNPs present among the *GmCKX* GFMs potentially affect CKX enzyme functioning. The observed *GmCKX* gene expression patterns and CK metabolite profiles supported the suggestion that the reduced *CKX* activity and increased CK accumulation, during soybean reproductive development enhanced yield parameters in SNP *GmCKX14* cultivars. The revealed natural genetic polymorphisms can be used as molecular indicators for identifying desirable soybean lines in larger genetic populations through marker assisted selection, which can save costs and time associated with breeding projects. Discovering useful genes, improving agricultural traits hidden in the plant genome, and applying these findings to crop breeding can pave the way for development of new, non-GMO soybean cultivars with higher yield and better abiotic stress resistance.

## Acknowledgements

Authors thank Ms. Amy Galer for analyzing soybean phytohormone data by HPLC-MS/MS.

