## Supplementary material for "The soybean (*Glycine max* L.) cytokinin oxidase/dehydrogenase multigene family; identification of natural variations for altered cytokinin content and seed yield": Table S

**Table S1.** Yield parameters of 16 soybean varieties used in this study in descending order. Highlighted in bold are the cultivars in which natural variation (SNP) was detected in *GmCKX14* gene.

| **Soybean** | **100 Seed weight (g)** | | |
| --- | --- | --- | --- |
|  | **>20** | **10-20** | **<10** |
| **Regular beans** | **IA1010LF**, OAC 06-14, **DH420**, **OAC Wallace**, OAC Bayfield | OAC Calypso, **DH748**, DH410SCN, RCAT Ruthven |  |
| **Natto beans** |  | OT08-13, Chikala | X5076-68B, X5076-159B, DH3604, X5076-108B, AC Colibri |
| **Soybean** | **Yield (t/ha)** | | |
|  | **>4** | **3-4** | **2-3** |
| **Regular beans** | **OAC Wallace**, OAC Calypso, DH410SCN, **DH748** | OAC Bayfield, OAC 06-14, RCAT Ruthven, **IA1010LF, DH420** |  |
| **Natto beans** |  | OT08-13, Chikala, X5076-159B | X5076-108B, DH3604, AC Colibri, X5076-68B |

**Table S2**. List of primers and PCR conditions used for amplification of soybean *CKX* genes.

| **Gene** | **PCR Fw Primer (5’-3’)** | **PCR Rw Primer (5’-3’)** | **Amplicon size (bp)** | **Annealing Tm** | **Extension time** |
| --- | --- | --- | --- | --- | --- |
| *GmCKX01* | TCCACCCCTTCACCTTTTCC | GTTCACAACCTCACAACACCCT | 2682 | 57^0^C | 2 m 40 s |
| *GmCKX02* | GCCTTGCAAAACAATATTCCACCC | ATGCCAGAGAGGTGCAGGAAG | 2753 | 52^0^C | 3 m 20 s |
| ^a^*GmCKX03* | TGAAAGAAAAATGCCACAGAGGA | AGCACATTATTTACTGGAAAATCCC | 4990 | 52^0^C | 5 m |
| ^a^*GmCKX04* | ATGGTAGCTGGGAAATACCCTTC | TTAATTGTTAAAGATTCTTTGT | 4328 | 55^0^C | 4 m 30 s |
| ^a^*GmCKX05* | CTCTTCCAGCCTCATCAAGT | CACATCCATGAAGTTTTGAGC | 4816 | 52^0^C | 5 m |
| ^a^*GmCKX06* | ATGGCTCTACACTACCCTTT | TTAAAACGCTGGCTGTAAT | 3975 | 50^0^C | 4 m |
| *GmCKX07a* | TGGTTGCTGAGAACTACCCT | CTCCCTCACTCCCAACAAGGA | 2197 | 53^0^C | 2 m 15 s |
| *GmCKX07b* | GCGGAAGCATCAAATGTCGTT | GCACACTTGTGATCAGCCAGCTA | 2175 | 52^0^C | 2 m 30 s |
| ^a^*GmCKX08* | ATGGCTCTAAACTACCCTTT | TTAAAACACTGGCTGTAATT | 4090 | 50^0^C | 5 m |
| *GmCKX09a* | CAAAGCAATAGTGCAACATTAACCA | TCGATTGTCTTACCGCAACCA | 2446 | 54^0^C | 2 m 30 s |
| ^a^*GmCKX09b* | TGCCATTTGGTAGTCTGGCA | AGCACCTGTACAACAAATTAGACG | 2396 | 55^0^C | 2 m 30 s |
| *GmCKX10a* | TCTACTGCGGCGTACTGGAC | TGACCAAGAAGAAAAAGCCCGA | 3295 | 56.5^0^C | 3 m 20 s |
| *GmCKX10b* | TGTACGCTTTTTATGTTATGCAGGA | TCCTTGTTGGGAGTGAGGGAG | 1975 | 56^0^C | 2 m |
| *GmCKX11a* | CGGAGGATGTGGCTAGGGTG | CGTGGGATTGGGTGTGTACG | 2114 | 59^0^C | 2 m 30 s |
| *GmCKX11b* | ATTCGAAGCATACCAACCGAAAT | AGCACTTGCTACTCAACATGAACC | 2123 | 57^0^C | 2 m 30 s |
| *GmCKX12* | ATGAGATACATCCTTGGAGAAC | TCATGAGAAGGTTATTGCTTT | 2993 | 54^0^C | 3 m |
| ^a^*GmCKX13* | ATGATGATTAAACTTCTCCA | TCATGAGAAGGTTATTGAT | 3297 | 50^0^C | 4 m |
| *GmCKX14* | ATGTCTTTCGTTCATGTTA | TACTTTGATTTGTTTACTGG | 2486 | 55^0^C | 2 m |
| ^a^*GmCKX15* | CGGCGATGCTAGCTTACCTGG | AAGGAGGTTGAGGTTGAGAGGT | 3407 | 57^0^C | 3 m 30 s |
| ^a^*GmCKX16*^b^ | CGCCAAAGCACTTCTCACAC | CTCTTATGAAGTCTTCCAAACTACG | 6148 | 53^0^C | 6 m 30 s |
| ^a^*GmCKX17* | ACTTGAACTTCGCCATAGCACG | ACAAGGGTTGAGGGATTCGGG | 7026 | 53^0^C | 7 m |

^a^ genes for which Expand Long Range dNTPack was used for amplification. ^b^ 6% DMSO was used in the PCR mixture.

**Table S3.** List of primers used for sequencing of soybean *CKX* genes using the Big Dye Terminator V3.1 Cycle sequencing kit.

| **Gene** | **Primer (5’ to 3’)** | **Direction** | **Amplicon sequenced (bp)^a^** |
| --- | --- | --- | --- |
| *GmCKX01 1* | CCTGGCCTTGAAGTGAATGG | Reverse | 1st exon (385-1) |
| *GmCKX01 2* | TTTGAAATGGGGTTTGCCTC | Forward | 1st exon (315-805) |
| *GmCKX01 3* | CTTTGGAGTGTGAATGGTGCA | Forward | 2-3 exons (888-1500) |
| *GmCKX01 4* | CCACTTCAAGCCAGCCAATT | Forward | 3-4 exons (1373-1930) |
| *GmCKX01 5* | GACAGTGCATATATTATTTTCTC | Forward | 5th exon (1955-2596) |
| *GmCKX02 1* | AGGCCTCCATGTACTTGTGC | Reverse | 1st exon (404-1) |
| *GmCKX02 2* | TCACTCACCATCAAGCATGT | Forward | 1st exon (294-854) |
| *GmCKX02 3* | TGGAGTGTGAATGGTGCATT | Forward | 2-3 exons (924-1554) |
| *GmCKX02 4* | TGGTGAAGCTGAAGCCATGA | Forward | 4th exon (1484-1984) |
| *GmCKX02 5* | TGAGGTAGATAGAATGTGGGGT | Forward | 5-7 exons (2138-2757) |
| *GmCKX03 1* | CTGCTATTGAGGTTAGTCAT | Reverse | 1st exon (400-1) |
| *GmCKX03 2* | TTCACTTCGAACAAGCGTGT | Reverse | 1st exon (656-400) |
| *GmCKX03 3* | AATTAGACCGGTCTATACAG | Reverse | 2nd exon (1828-2318) |
| *GmCKX03 4* | ACGTCTAACTTGACCTAACA | Forward | 4th exon (2959-3449) |
| *GmCKX03 5* | TCACTCAAATTGTATGGAACCCA | Forward | 5th exon (3458-3948) |
| *GmCKX03 6* | TCCATTGGGCCTGACAGTAG | Forward | 6th exon (4452-4934) |
| *GmCKX04 1* | CCCTCGTCGAGTGCCCTT | Reverse | 1st exon (374-1) |
| *GmCKX04 2* | CATCGCGAGGCTGATCAAG | Forward | 1-2 exons (267-826) |
| *GmCKX04 3* | AACGGGCTGCTGAATTTTAA | Forward | 3rd exon (1640-2060) |
| *GmCKX04 4* | GGGACTCATCGCATAAGCCA | Forward | 4th exon (2381-2941) |
| *GmCKX04 5* | GTAGTATATAGTTTTCGTACC | Reverse | 5th exon (2771-3401) |
| *GmCKX04 6* | AGGTCATTTCGTACCAATTA | Forward | 5-6 exons (3389-4028) |
| *GmCKX05 1* | CTCTTCCAGCCTCATCAAGT | Forward | 1^st^ exon (1-550) |
| *GmCKX05 2* | CATTGGGTACACTTGATACA | Forward | 2-3 exons (2977-3680) |
| *GmCKX05 3* | AGTTGTCTACATTGTCATCTA | Forward | 4^th^ exon (3724-4254) |
| *GmCKX05 4* | CACATCCATGAAGTTTTGAGC | Reverse | 5^th^ exon (4816-4300) |
| *GmCKX06 1* | TCGTCTATAGAAGATGGAC | Reverse | 1st exon (450-1) |
| *GmCKX06 2* | CAAGATTTATGATCTTATCCTAC | Reverse | 1st exon (727-450) |
| *GmCKX06 3* | ACTAGCTAGGTCCACGGTCA | Forward | 2nd exon (1338-1828) |
| *GmCKX06 4* | TGGGACTAGTTGTGAGTGTGC | Forward | 3rd exon (2250-2800) |
| *GmCKX06 5* | TGGACAAGGTACCCTCGTCT | Forward | 4th exon (2672-3232) |
| *GmCKX06 6* | AGGAGTGAAGAGTGTGATGCA | Forward | 5th exon (3149-3709) |
| *GmCKX07a1* | TCGCCGCAATCTTGAAGG | Reverse | 1st exon (400-1) |
| *GmCKX07a2* | AACTATTGGATATTGGAAGC | Reverse | 1st exon (800-400) |
| *GmCKX07a3* | CGCAAAGCTTGGTGGTTGAA | Reverse | 2nd exon (1910-1410) |
| *GmCKX07b1* | GATGGGACTCGTTGCATAAG | Forward | 3rd exon (2170-2729) |
| *GmCKX07b2* | GTGGTATATATAGTTTTCGTACC | Reverse | 4th exon (3600-2967) |
| *GmCKX07b3* | TTGTGATCAGCCAGCTAGCT | Reverse | 5th exon (4155-3521) |
| *GmCKX08 1* | AGACACTAATCGCAACCCCA | Reverse | 1st exon (406-1) |
| *GmCKX08 2* | CATGGCTCGTGATGGGATTG | Forward | 1st exon (332-822) |
| *GmCKX08 3* | CGATGCAAACTTTCCAAGGG | Forward | 2nd exon (1354-1843) |
| *GmCKX08 4* | TGGGACTGGTTGCACATACA | Forward | 3rd exon (2437-2997) |
| *GmCKX08 5* | TGTGGACAAGGTAACCTCGT | Forward | 4th exon (2730-3290) |
| *GmCKX08 6* | CAGTCTTGGTTTATCCCATGA | Forward | 5th exon (3285-3915) |
| *GmCKX09a1* | GGCTAGGGAAGAGAATAACC | Reverse | 1st exon (605-1) |
| *GmCKX09b1* | AGCGATTAAGCGTACATGAT | Reverse | 2-3 exons (3212-2761) |
| *GmCKX09b2* | AATCTTCATTAGTTATGACCCTT | Forward | 3-4 exons (3065-3694) |
| *GmCKX09b3* | GAATGCTAGATCTTCTTGGT | Forward | 4-5 exons (3517-4146) |
| *GmCKX09b4* | TGGTATGTCTGTGGAGTTAATACTT | Forward | 5-6 exons (4036-4735) |
| *GmCKX10a1* | TCTACTGCGGCGTACTGGAC | Forward | 1st exon (1-550) |
| *GmCKX10a2* | GCATGTTGCGTACAGAATCT | Forward | 1st exon (574- 1200) |
| *GmCKX10a3* | ATTCGATAAACGGGCAAGCC | Forward | 1st exon (1150-1670) |
| *GmCKX10a4* | TAGTTGTCCTTGTTCCCCAA | Forward | 2nd exon (1681-2100) |
| *GmCKX10a5* | AGATTAACGGATAATTGGATG | Forward | 3rd exon (2306-3075) |
| *GmCKX10b1* | AGACCTAAACACTGACACTCGA | Reverse | 4th exon (5841-5378) |
| *GmCKX10b2* | AGTGACAAGTCAAAGGATTG | Forward | 5th exon (6650-7275) |
| *GmCKX11a1* | GTAAAAATTAACCGTGAATGCA | Reverse | 1st exon (6-560) |
| *GmCKX11a2* | TGTTCCCCAAGAGGAAAAGT | Forward | 2nd exon (719-1138) |
| *GmCKX11a3* | ATCATACATCATCGTGGTGGTA | Forward | 3rd exon (2130-1622) |
| *GmCKX11b1* | TCGAAGCATACCAACCGAAA | Forward | 4th exon (4330-4820) |
| *GmCKX11b2* | ATGGTTTCAAGTGTGCAGTA | Forward | 5th exon (5886-6446) |
| *GmCKX12 1* | TGTGGATTTGATTGCCTAGC | Reverse | 1-2 exons (500-1) |
| *GmCKX12 2* | GCTAGGCAATCAAATCCACA | Forward | 1-3 exons (500-950) |
| *GmCKX12 3* | AACAGGGGAGGTGTTGAACT | Forward | 3-4 exons (896-1426) |
| *GmCKX12 4* | GCGACTGCGACCTATATTTA | Forward | 5th exon (1580-2140) |
| *GmCKX12 5* | TTGTTTGCAGGTGGGACAAC | Forward | 6th exon (2040-2530) |
| *GmCKX13 1* | CCTCCTGAGACATCAACATA | Reverse | 1st exon (467-1) |
| *GmCKX13 2* | GGCCCATGGAGGAGTTGTTA | Forward | 1-2 exons (410-1050) |
| *GmCKX13 3* | TGAGGAGCAGAATGGTGAACT | Forward | 2-3 exons (964-1524) |
| *GmCKX13 4* | TCGACACTTGATGGTAGAAATCA | Forward | 4th exon (1631-2240) |
| *GmCKX13 5* | TGCCAGTACATGTCCTATTT | Forward | 5th exon (2568-2997) |
| *GmCKX14 1* | GAATGGCTATGCAACTAAGG | Reverse | 1-3 exons (538-1) |
| *GmCKX14 2* | GAGATACCATTACCCACCTCGT | Forward | 3rd exon (440-1000) |
| *GmCKX14 3* | ACTCTAAGGTACGGGTTGGC | Forward | 3-5 exons (933-1600) |
| *GmCKX14 4* | AGCACCTGCCATGGTAATGA | Forward | 4-6 exons (1418-2000) |
| *GmCKX14 5* | ACGTGGAAGAAATCGATGCAG | Forward | 5-6 exons (1768-2328) |
| *GmCKX15 1* | CCAATTCCGTTACGTTCGAT | Reverse | 1-3 exons (478-1) |
| *GmCKX15 2* | CGAGGTCGTGGACGGATTAT | Forward | 3-4 exons (440-942) |
| *GmCKX15 3* | AGAGTCGCTGGTGGAGGAAT | Forward | 4th exon (640-1260) |
| *GmCKX15 4* | TGCCCTCGTTAAGTTTAGCA | Forward | 5th exon (1542-2102) |
| *GmCKX15 5* | AGTCATGCGCCTTGAATTGA | Forward | 6th exon (2878-3418) |
| *GmCKX16 1* | TTGCGATTCGGAGCAGACTA | Reverse | 1st exon (584-1) |
| *GmCKX16 2* | CGTCAGCGGCCAGTCCTT | Forward | 1-2 exons (528-1160) |
| *GmCKX16 3* | TAGGAAAATATGTCAATGTCG | Forward | 3rd exon (1696-2300) |
| *GmCKX16 4* | AGTAATGTAGTCATAAGGACC | Forward | 4th exon (5280-5700) |
| *GmCKX17 1* | ACTTGAACTTCGCCATAGCACG | Forward | 1st exon (1-500) |
| *GmCKX17 2* | CGCAAACCGCCAACGTAA | Forward | 1-2 exons (509-1160) |
| *GmCKX17 3* | CTCCTCCTCCTCCTTCCAGT | Forward | 3rd exon (1927-2550) |
| *GmCKX17 4* | CAATTGTGTCTGCCTCAAACA | Forward | 4th exon (6388-6948) |

^a^ The nucleotide number of the sequenced region represents its position from the beginning of amplicon using the PCR primers listed in this table.

**Table S4**. List of PCR and sequencing primers used for re-sequencing analysis to confirm the presence of polymorphisms in sequences of soybean *CKX* genes.

| **Gene** | **PCR primers (5`-3`)** | **Sequencing primers (5`-3`)** | **Amplicon sequenced^a^** |
| --- | --- | --- | --- |
| *GmCKX04-1* | ATTCGAAGCATACCAACCGAAAT  AGCACTTGCTACTCAACATGAACC | Fw-ATGGTTTCAAGTGTGCAGTA  Rw-AGCACTTGCTACTCAACATGAACC | 5th exon- (1678-2053)  5th exon- (2123-1700) |
| *GmCKX04-2* | CGGCGATGCTAGCTTACCTGG,  AAGGAGGTTGAGGTTGAGAGGT | Fw- CGGCGATGCTAGCTTACCTGG  Rw- CCAATTCCGTTACGTTCGAT | 1st exon- (1-520)  1st exon- (478-1) |
| *GmCKX06* | TCTACTGCGGCGTACTGGAC  TGACCAAGAAGAAAAAGCCCGA | Fw-GCATGTTGCGTACAGAATCT  Rw-GGCTTGCCCGTTTATCGAAT | 1st exon- (574-1205)  1st exon- (1116-696) |
| *GmCKX09-1* | ATGAGATACATCCTTGGAGAAC | Fw-ATCCTTCAACCCACGAGACC  Rw-AGTCTCAATTCGGGTCGCAA  Fw-ACTTCTGTTGTGATTCCAGAGGA  Rw-TCATGAGAAGGTTATTGCTTT | 4th exon- (1270-1550)  4th exon- (1520-1210)  6th exon- (2063-2413)  6th exon- (2376-2000) |
|  | TCATGAGAAGGTTATTGCTTT |  |  |
| *GmCKX14* | ACCGAAACGAAATCCACTCT  ATGTCAAGGTCAAGCTCACA | Fw AGCGTACCTAGAACACTTCGTG  Rw- TTGCGATTCGGAGCAGACTA | 1^st^ exon- (29-612)  1^st^ exon- (593-1) |

^a^ The nucleotide number of the sequenced region represents its position from the beginning of amplicon using the PCR primers listed in this table.

**Table S5**. List of primers used in qRT-PCR analysis of soybean *CKX* gene expression in seeds of OAC Wallace.

| **Gene** | **PCR Fw Primer (5’-3’)** | **PCR Rw Primer (5’-3’)** |
| --- | --- | --- |
| *GmCKX03* | TGGGCAAGCCTTCAAACATGG | AAGGTCAGCGTTTCGGTTCC |
| *GmCKX04-1* | ACAAAAACAAGTGGGACCAGCG | AGCGTCTCAGTATCCAATGCCG |
| *GmCKX04-2* | TATGCTAGGGATGCAGAGTCGC | CATCCATTGGCCCGGTTATCAC |
| *GmCKX06* | AGCAGCAAGGGAGAGATTCGAC | AGCTGCAAAGAACGAAGACCTC |
| *GmCKX09-1* | CCCGGTCCAAGCTAGTCATTTC | TGCTGCATCGATTTCTTCCACG |
| *GmCKX09-2* | TGGACTTGCACCTATGTCCTGG | AGGACCATAGCGGAATGTCTGG |
| *GmCKX09-3* | TCCCTTGAACGTGGACTCACAC | ATGCCACCCATGCCAGCATTAG |
| *GmCKX11* | GGTCAAGTTGCGTTCAAAAGGC | AGACAACCTCTGCGAAATGGTG |
| *GmCKX12* | CAAGACCCGGTTCAAGCTAGTC | TGCTGCATCGATTTCTTCCACG |
| *GmCKX13* | AACTTGGAGTTATTCCACGCGG | ACTTAACCCTTTTGGGGGCTGG |
| *GmCKX14* | CGTAACCAAGACCATCCTTCCG | CACGCAGAACTTTAGACCCTCG |
| *GmCKX15* | TGGACTTGCACCTATGTCCTGG | AGGACCATAGCGGAATGTCTGG |
| *GmCKX17-1* | AGTTATTCCACGCGGTTCTTGG | CCCACTTAACCCTTTTGGGTGC |
| *GmCKX17-2* | ACACTTGAGCGTGGACTCACAC | GCCACTTATCCCAGCATTGGAG |
| *GmCKX17-3* | GGCGAAACCTTAGTCTGTTCCG | CGCGAGTTATGATGCCGAACTG |
| *GmCKX19* | TGGGCAAGCCTTCAAACATGG | AGGTCAGCATTTCGGTTCCCTG |

**Table S6a.** Cytokinin [pmol/gFW] in pods (R4 & R5 stages) of 4 soybean cultivars analysed using HPLC-(ESI+)-MS/MS. Data are means of 3 biological replicates (n=3±SE).

| Cultivar | | OAC 06-14 | DH 420 | OAC Wallace | DH 3604 | OAC 06-14 | DH 420 | OAC Wallace | DH 3604 |
| --- | --- | --- | --- | --- | --- | --- | --- | --- | --- |
|  | | Stage | | | | | | | |
| Cytokinin [pmol/gFW] | | R4 (pod) | | | | R5 (pod) | | | |
| Free bases (FB) | tZ | 53.07±3.07 | 59.63±9.91 | 45.84±6.85 | 52.94±7.35 | 40.97±8.20 | 40.41±7.06 | 35.39±6.74 | 16.20±9.35 |
|  | cZ | 38.37±19.99 | 8.62±0.69 | 20.39±6.85 | 5.01±2.14 | 95.35±27.28 | 10.08±2.28 | 24.74±4.82 | 7.96±3.41 |
|  | DZ | 26.70±1.78 | 31.11±8.34 | 21.26±0.28 | 19.12±6.80 | 23.98±2.67 | 27.10±2.08 | 20.14±5.32 | 10.50±2.44 |
|  | iP | 0.87±0.04 | 0.57±0.09 | 1.16±0.08 | 2.23±0.19 | 1.15±0.24 | 0.68±0.11 | 0.83±0.09 | 1.43±0.43 |
| Ribosides (RB) | tZR | 20.55±2.12 | 29.24±2.70 | 18.20±3.01 | 80.41±24.00 | 29.27±3.57 | 29.22±11.38 | 29.22±6.69 | 20.60±4.83 |
|  | cZR | 41.63±10.23 | 56.91±21.50 | 41.65±4.40 | 44.99±18.94 | 73.22±3.86 | 103.71±21.96 | 39.03±9.97 | 89.60±28.19 |
|  | DZR | 74.00±11.77 | 35.71±1.60 | 35.13±8.12 | 57.65±9.47 | 30.16±3.11 | 23.33±3.23 | 10.40±3.07 | 13.18±2.62 |
|  | iPR | 2.34±0.06 | 1.83±0.40 | 3.09±0.49 | 3.26±0.60 | 3.19±0.59 | 1.91±0.32 | 1.89±0.16 | 2.88±0.17 |
| Nucleotides (NT) | tZNT | 3.85±2.14 | 8.74±2.31 | 0.89±0.14 | 2.73±0.80 | 2.39±0.09 | 6.48±1.20 | 1.85±0.39 | 1.94±0.56 |
|  | cZNT | 25.57±2.16 | 210.63±71.52 | 19.53±2.70 | 28.64±4.14 | 45.64±2.75 | 72.17±18.34 | 26.76±6.62 | 41.95±8.89 |
|  | DZNT | 0.73±0.18 | 1.62±0.44 | 0.21±0.04 | 0.52±0.15 | 0.41±0.04 | 0.71±0.09 | 0.08±0.03 | 0.27±0.14 |
|  | iPNT | 1.04±0.47 | 14.06±5.21 | n.d. | 0.80±0.12 | 1.40±0.57 | 1.07±0.23 | n.d. | 1.03±0.34 |
| Methylthiols (MET) | 2MeSZ | 5.20±1.01 | 5.36±1.16 | 10.74±0.31 | 75.59±49.32 | 36.27±24.15 | 6.08±1.46 | 9.76±0.33 | 21.33±10.81 |
|  | 2MeSiP | 0.20±0.06 | 0.35±0.06 | 0.41±0.03 | 3.24±1.73 | 2.84±2.03 | 0.37±0.11 | 0.44±0.04 | 1.52±0.80 |
|  | 2MeSZR | 2.76±1.26 | 5.35±2.41 | 4.87±0.68 | 6.04±1.59 | 158.09±126.04 | 17.63±8.74 | 4.63±1.12 | 12.70±3.65 |
|  | 2MeSiPR | n.d. | n.d. | n.d. | 1.85±1.51 | n.d. | n.d. | n.d. | n.d. |
| Glucosides  (GLUC) | DZOG | 3.00±0.30 | 4.15±1.22 | 4.22±0.64 | 2.81±0.90 | 7.92±1.41 | 9.84±2.59 | 6.52±1.29 | 5.76±0.74 |
|  | DZ9G | n.d. | n.d. | n.d. | n.d. | n.d. | n.d. | n.d. | n.d. |
|  | tZOG | 1.90±0.18 | 0.96±0.11 | 2.63±0.19 | 3.59±1.71 | 1.66±0.26 | 1.16±0.20 | 2.12±0.30 | 24.64±5.10 |
|  | cZOG | n.d. | n.d. | n.d. | n.d. | n.d. | n.d. | n.d. | n.d. |
|  | tZ9G | n.d. | n.d. | n.d. | n.d. | n.d. | n.d. | n.d. | n.d. |
|  | cZ9G | n.d. | n.d. | n.d. | n.d. | n.d. | n.d. | n.d. | n.d. |
|  | DZROG | 43.47±8.55 | 33.77±1.65 | 98.87±4.73 | 160.82±50.48 | 96.80±22.86 | 95.61±32.66 | 170.78±95.99 | 64.05±25.21 |
|  | tZROG | 3.27±0.37 | 1.55±0.09 | 2.96±0.16 | 6.40±1.55 | 3.09±0.37 | 1.42±0.32 | 2.87±0.88 | 2.33±0.70 |
|  | cZROG | 2.64±0.26 | 2.86±0.30 | 3.32±0.30 | 7.50±2.44 | 1.28±0.33 | 0.66±0.01 | 1.74±0.17 | 3.63±1.13 |

n.d. – not detected

**Table S6b**. Cytokinin [pmol/gFW] in seeds (R5.5 & R6 stages) of 4 soybean cultivars analysed using HPLC-(ESI+)-MS/MS. Data are means of 3 biological replicates (n=3±SE).

| Cultivar | | OAC 06-14 | DH 420 | OAC Wallace | DH 3604 | OAC 06-14 | DH 420 | OAC Wallace | DH 3604 |
| --- | --- | --- | --- | --- | --- | --- | --- | --- | --- |
|  | | Stage | | | | | | | |
| Cytokinin [pmol/gFW] | | R5.5 (seed) | | | | R6 (seed) | | | |
| Free bases (FB) | tZ | 8.51±1.36 | n.d. | 31.47±9.35 | n.d. | 1.36±0.64 | n.d. | 23.55±1.63 | 0.47±0.38 |
|  | cZ | 424.21±130.17 | 246.47±23.30 | 118.05±24.49 | 116.99±12.31 | 2.26±1.84 | 63.72±11.02 | 138.06±17.55 | 64.86±9.53 |
|  | DZ | 7.33±3.08 | 7.49±0.48 | 13.33±2.60 | n.d. | n.d. | n.d. | 4.62±1.92 | n.d. |
|  | iP | n.d. | n.d. | n.d. | n.d. | n.d. | n.d. | n.d. | n.d. |
| Ribosides (RB) | tZR | 18.14±5.19 | 11.13±1.09 | 12.76±2.54 | 5.65±0.51 | 3.02±0.54 | 1.24±0.08 | 6.45±1.26 | 1.94±0.46 |
|  | cZR | 18.86±4.13 | 25.05±1.99 | 9.52±3.08 | 22.62±0.39 | 3.37±0.26 | 5.01±0.46 | 2.79±0.33 | 18.79±2.16 |
|  | DZR | 15.59±1.29 | 7.54±0.76 | 5.09±0.53 | 6.67±0.56 | 21.03±2.70 | 2.46±0.11 | 3.68±0.24 | 5.74±0.68 |
|  | iPR | 16.33±1.35 | 8.39±0.85 | 5.33±0.55 | 7.19±0.60 | 22.04±2.83 | 2.76±0.13 | 3.87±0.25 | 6.23±0.73 |
| Nucleotides (NT) | tZNT | 3.52±1.50 | 1.82±1.04 | 16.98±2.65 | 5.29±1.26 | 11.01±4.62 | 3.24±1.98 | 8.00±3.08 | 5.18±4.23 |
|  | cZNT | 31.89±12.80 | 32.05±7.11 | 43.44±13.62 | 44.48±1.86 | 36.77±21.12 | 9.85±1.29 | 18.38±3.63 | 41.53±13.30 |
|  | DZNT | 40.47±5.73 | 24.42±10.37 | 34.68±9.23 | 28.46±6.64 | 27.25±3.84 | 24.40±2.27 | 38.78±11.24 | 47.68±7.62 |
|  | iPNT | 9.52±3.90 | 4.15±1.70 | 25.27±8.12 | 7.09±1.78 | 17.48±4.05 | 7.21±1.53 | 12.47±0.90 | 9.50±3.89 |
| Methylthiols (MET) | 2MeSZ | n.d. | n.d. | n.d. | n.d. | n.d. | n.d. | n.d. | n.d. |
|  | 2MeSiP | n.d. | n.d. | n.d. | n.d. | n.d. | n.d. | n.d. | n.d. |
|  | 2MeSZR | n.d. | n.d. | n.d. | n.d. | n.d. | n.d. | n.d. | n.d. |
|  | 2MeSiPR | n.d. | n.d. | n.d. | n.d. | n.d. | n.d. | n.d. | n.d. |
| Glucosides  (GLUC) | DZOG | 13.17±3.43 | 8.94±2.90 | 4.76±0.74 | 6.85±0.49 | 3.32±0.15 | 6.05±1.51 | 1.34±0.65 | 7.90±1.64 |
|  | DZ9G | 15.35±6.58 | 8.57±0.86 | 10.48±1.16 | 11.30±0.98 | 10.10±1.19 | 5.88±0.18 | 15.51±3.14 | 7.39±1.09 |
|  | tZOG | n.d. | n.d. | n.d. | n.d. | n.d. | n.d. | n.d. | n.d. |
|  | cZOG | n.d. | n.d. | n.d. | n.d. | n.d. | n.d. | n.d. | n.d. |
|  | tZ9G | 10.83±2.14 | 11.81±1.73 | 4.44±0.33 | 9.20±0.79 | 3.22±0.23 | 2.57±1.14 | 2.50±0.54 | 11.64±3.23 |
|  | cZ9G | 5.91±0.65 | 16.53±3.52 | 3.23±0.39 | 36.53±3.14 | n.d. | 6.31±1.44 | 2.69±0.63 | 15.06±3.19 |
|  | DZROG | 104.40±39.55 | 43.99±18.62 | 65.19±19.76 | 10.34±3.41 | n.d. | 24.23±19.78 | 1.68±1.38 | 7.48±6.10 |
|  | tZROG | 12.38±5.09 | 0.32±0.15 | 5.42±1.61 | n.d. | n.d. | 0.10±0.08 | n.d. | n.d. |
|  | cZROG | n.d. | n.d. | n.d. | n.d. | n.d. | n.d. | n.d. | n.d. |

n.d. – not detected

**I**

**III**

**IV**

**V**

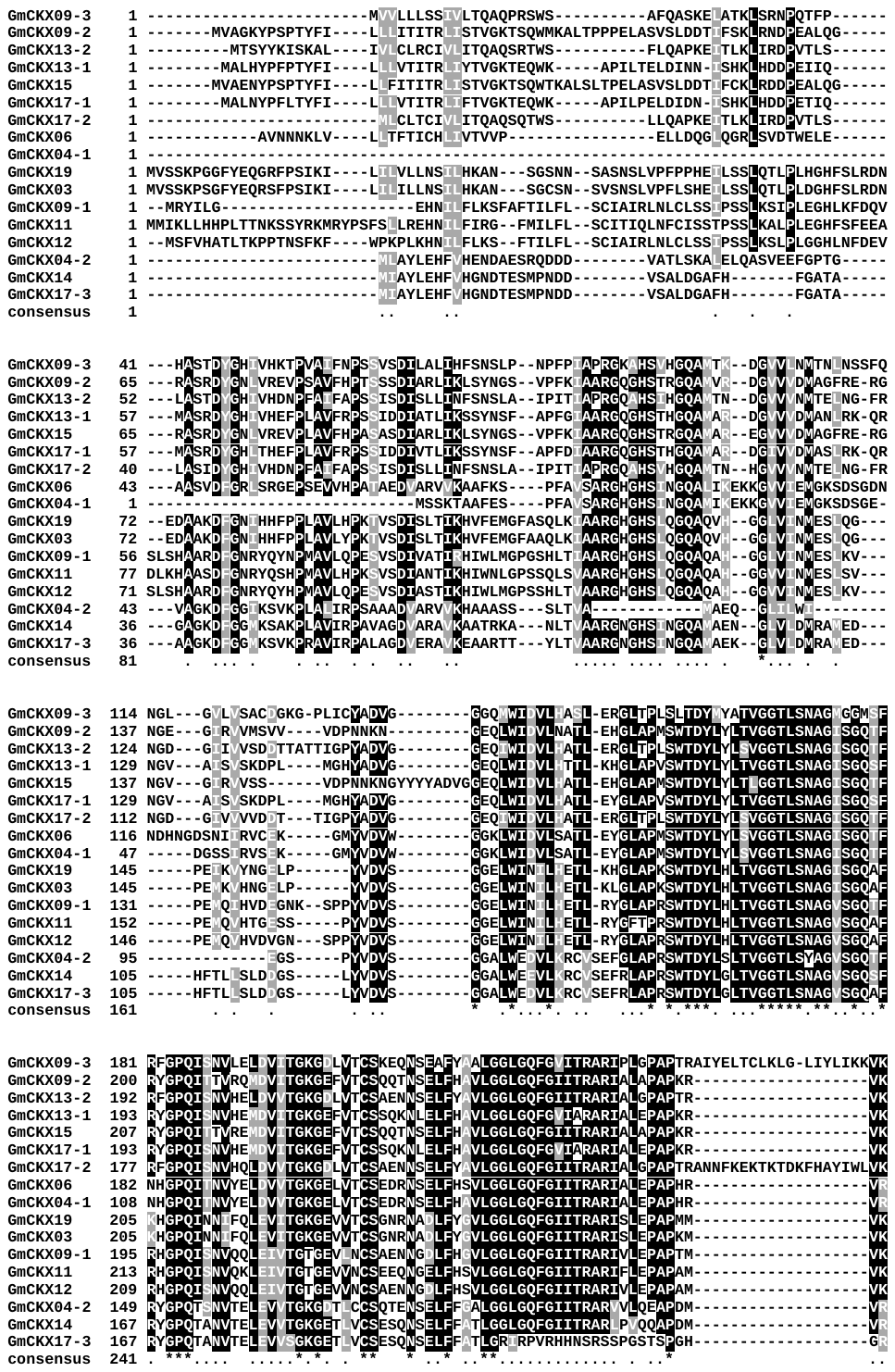

**I**

**III**

**IV**

**V**

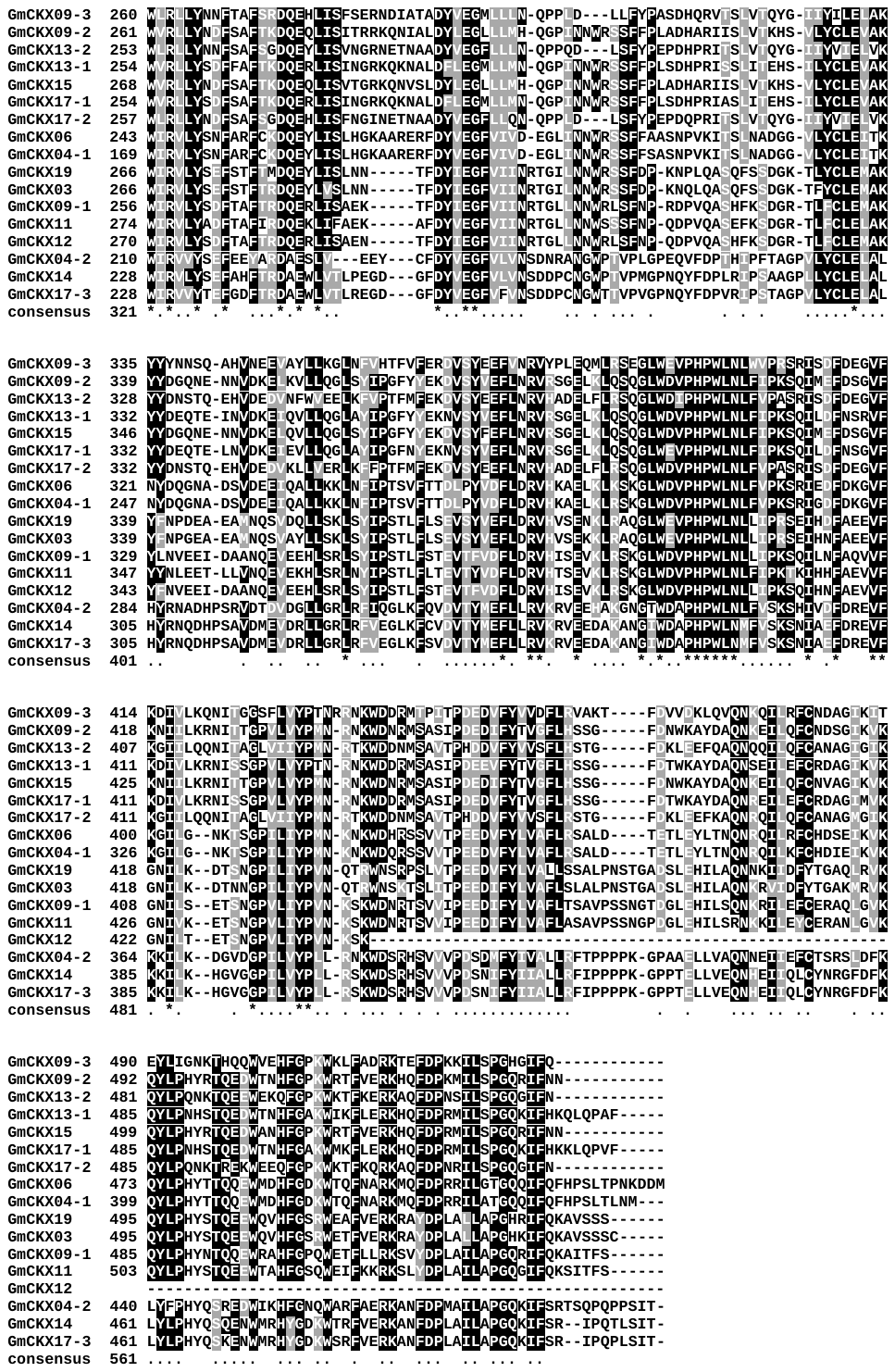

**Fig. S1** Boxshade representation (with >70% similarity) of the 17 soybean CKX protein sequences aligned using the ClustalW interface in MEGA software (Penn State University, USA). Identical amino acid residues are highlighted in black and similar amino acid residues are highlighted in grey. A consensus sequence with “*” representing identity and “.” representing similarity is also presented. The phylogenetic clades, to which each of the CKX proteins belonged to, are presented to the left on the top panel
